# Cooperative Binding Mitigates the High-Dose Hook Effect

**DOI:** 10.1101/021717

**Authors:** Ranjita Dutta Roy, Christian Rosenmund, Melanie I Stefan

**Affiliations:** Department of Medicine Solna, Karolinska Institutet, Stockholm, Sweden.; NWFZ, Charité Crossover, Charite Universitätsmedizin, Berlin, Germany.; Department of Neurobiology, Harvard Medical School, Boston, United States.; Babraham Institute, Cambridge, United Kingdom.; Centre for Integrative Physiology, University of Edinburgh, Edinburgh, United Kingdom.

**Keywords:** prozone effect, high-dose Hook effect, mechanistic model, cooperativity, allostery, calmodulin

## Abstract

**Background:** The high-dose hook effect (also called prozone effect) refers to the observation that if a multivalent protein acts as a linker between two parts of a protein complex, then increasing the amount of linker protein in the mixture does not always increase the amount of fully formed complex. On the contrary, at a high enough concentration range the amount of fully formed complex actually decreases. It has been observed that allosterically regulated proteins seem less susceptible to this effect. The aim of this study was two-fold: First, to investigate the mathematical basis of how allostery mitigates the prozone effect. And second, to explore the consequences of allostery and the high-dose hook effect using the example of calmodulin, a calcium-sensing protein that regulates the switch between long-term potentiation and long-term depression in neurons.

**Results:** We use a combinatorial model of a “perfect linker protein” (with infinite binding affinity) to mathematically describe the hook effect and its behaviour under allosteric conditions. We show that allosteric regulation does indeed mitigate the high-dose hook effect. We then turn to calmodulin as a real-life example of an allosteric protein. Using kinetic simulations, we show that calmodulin is indeed subject to a hook effect. We also show that this effect is stronger in the presence of the allosteric activator Ca^2+^/calmodulin-dependent kinase II (CaMKII), because it reduces the overall cooperativity of the calcium-calmodulin system. It follows that, surprisingly, there are conditions where increased amount of allosteric activator actually *decrease* the activity of a protein.

**Conclusions:** We show that cooperative binding can indeed act as a protective mechanism against the hook effect. This will have implications *in vivo* where the extent of cooperativity of a protein can be modulated, for instance, by allosteric activators or inhibitors. This can result in counterintuitive effects of decreased activity with increased concentrations of both the allosteric protein itself and its allosteric activators.

## Background

Since the early 20^th^ century, immunologists have noted that more is not always better: Increasing the amount of antibody in an antibody-antigen reaction could reduce, instead of increase, the amount of precipitating antibody-antigen complex [1]. Similarly, mice receiving larger doses of anti-pneumococcus horse serum were not more, but less protected against pneumococcus infection [2, 3]. There was clearly a range of antibody concentrations above the optimum at which no effects (or negative effects) were obtained. This region of antibody concentrations was named the prozone, and the related observation the “prozone effect” [1, 2, 3] or (after the shape of the complex formation curve) the “high-dose hook effect” (reviewed in [4, 5]).

Over the following decades, the high-dose hook effect became better understood beyond its first application in immunology, and as a more general property of systems involving multivalent proteins. In 1997, Bray and Lay showed using simulations of various types of protein complexes that the prozone effect is a general phenomenon in biochemical complex formation, and occurs whenever one protein acts as a “linker” or “bridge” between parts of a complex [6]. This was corroborated using a mathematical model of an antibody with two antigen-binding sites by Bobrovnik [7] and in a DNA-binding experiment by Ha et al. [8].

The hook effect thus results from partially bound forms of the “linker” proteins competing with each other for binding partners, and as a consequence, there is a regime of concentrations where adding more linker protein will decrease the amount of fully formed complexes, rather than increase it (see figure 1).

**Figure 1.**
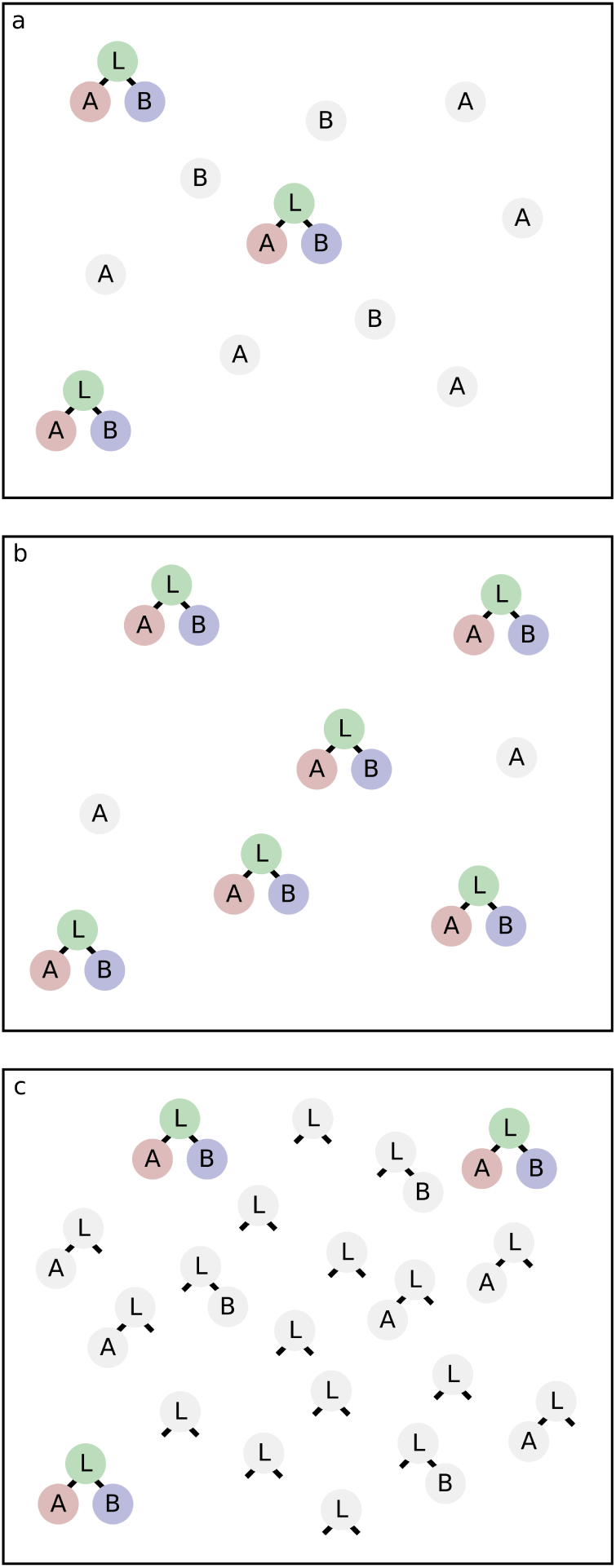
Binding of ligands A, B to a bivalent linker protein L. a) Low linker concentration: availability of L limits the formation of total complexes (LAB, in colour). b) Linker concentration on the order of ligand concentration: Formation of fully formed complex (LAB) reaches its maximum. c) Concentration of linker L much higher than that of A or B: partially bound forms prevail, and formation of fully formed complex (LAB) goes down in absolute terms.

Are all complexes with a central multivalent “linker” protein equally susceptible to the hook effect? Based on simulation of allosterically regulated proteins using the Allosteric Network Compiler (ANC), Ollivier and colleagues suggested that allostery can mitigate the prozone effect [9].

In this case, ligand binding to the linker protein is cooperative (reviewed in [10]), and the simulations by Ollivier et al. showed that the higher the cooperativity, the less pronounced the hook effect [9].

This agrees with what we know about cooperative binding: If ligand binding to one site is conducive to ligand binding to other sites, this will favour the formation of fully assembled complex over partial complexes, and thus increase the total amount of fully formed complex at a given linker concentration, compared to the non-cooperative case. In other words, partially bound forms of the linker protein still compete among themselves for binding partner, but cooperative binding skews the competition in favour of the forms that have more binding sites occupied and are thus closer to the fully bound form.

In this paper, we formalise and further develop these ideas. We first provide a mathematical description of the principle behind the high-dose hook effect and show that it is indeed smaller for proteins that display cooperative ligand binding.

We then go on to examine how this applies to allosteric proteins. We have decided to investigate the case of calmodulin, an allosteric tetra-valent calcium binding protein that is present in many tissues of the human body. In neurons, calmodulin acts as a switch between long-term potentiation and long-term depression of a synaptic connection in response to the frequency, duration and amplitude of a calcium signal [11]. We investigate the effects of both the hook effect itself and the allosteric nature of calmodulin under conditions comparable to its cellular environment.

## Results

### A combinatorial model shows that increasing amounts of linker protein lead to decreasing amounts of complex

We start by looking at a case in which a linker protein L binds perfectly (i.e. with an infinitely small *K_d_*) to one molecule each of A and B to form a ternary complex (LAB, see figure 1). The binding sites for A and B are separate and have the same affinity for the linker L.

In the following, we will denote amounts or numbers of molecules with lower-case letters: *a* will be the number of molecules of A, *b* the number of molecules of B, and *λ* the number of molecules of L. Without loss of generality, we will assume that *b* ≤ *a*.

In this case (see Methods section for details), we can write the expected amount of LAB as a function of *λ* as a three-part function:

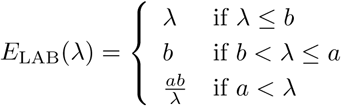

A plot of the above function for a = 80, b=50, and *λ* = 1 to 400 is shown in figure 2 (black line). In order to visualise the stochastic fluctuation around those expected values, for each value of *λ*, the figure also shows the result of 100 stochastic simulations (grey dots). For each of these, *a* molecules of L were randomly chosen for binding to A, and *b* molecules of L were randomly chosen for binding to B, and we then counted the resulting number of molecules of L that were bound to both A and B (see “Methods”).

**Figure 2.**
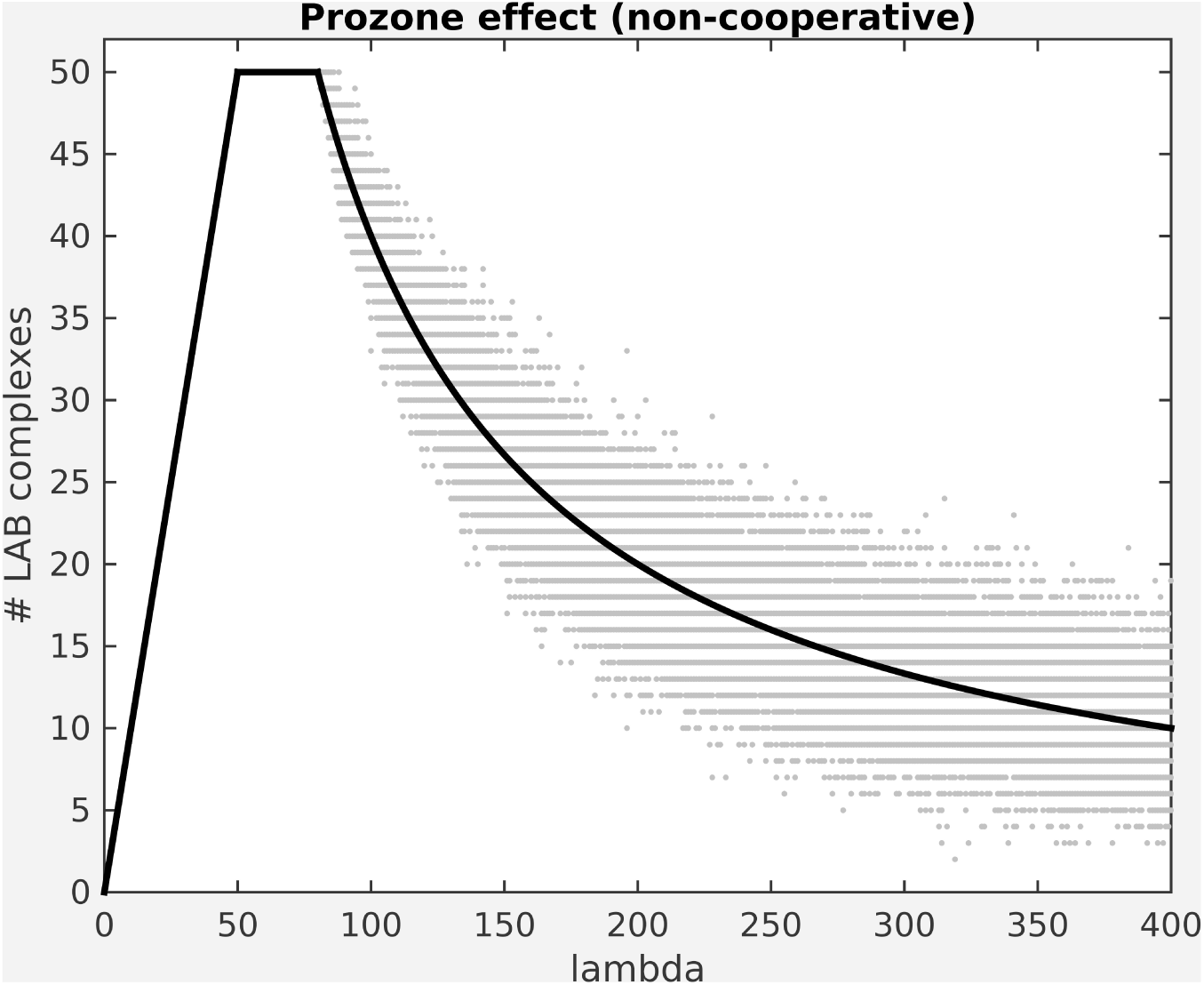
Prozone effect for a Linker protein without cooperativity, assuming perfect binding. Black line: Expected value. Grey dots: Results of 100 stochastic simulations. Amount of linker protein (lambda) varied from 1 to 400, amounts of proteins A and B were 80 and 50, respectively. Simulations were run using MATLAB[36].

**Figure 3.**
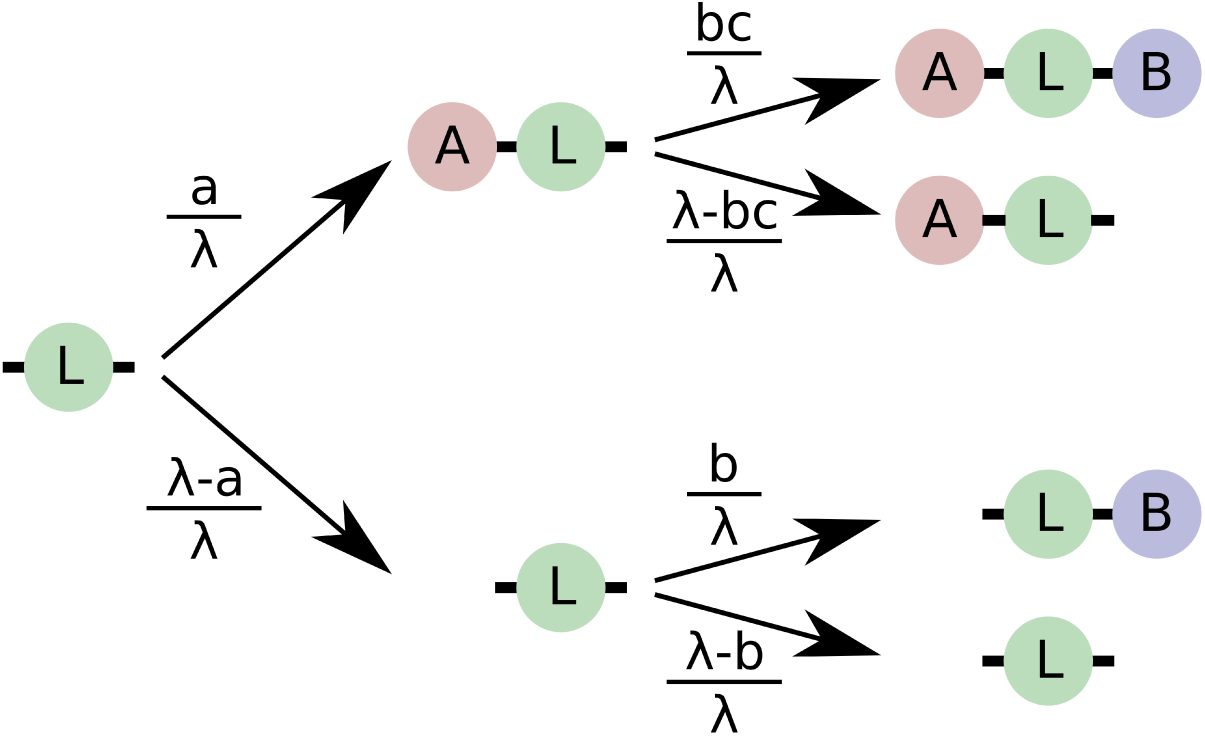
Probability of binding events for a cooperative linker L when both *a* and *b* are smaller than *λ*. For each L, the arrows are marked with the probabilities of the associated binding event. The amount of cooperativity is indicated by a multiplicative factor *c*, where *c >* 1 denotes positive cooperativity, and *c* = 1 in the absence of cooperativity.

As we can see, the amount of fully bound complex will first increase with increasing amounts of L, then stay constant (at *b*) until the amount of L exceeds the amounts of both A and B, and then go down again as L increases further. In other words, for large enough L, adding L will decrease the expected amounts of fully bound complex LAB. This is the high-dose hook effect.

### Cooperative binding attenuates the high-dose hook effect

Now, how does the situation change if binding to L is cooperative, i.e. if binding of L to a molecule of A (or B) is more likely when a molecule of B (or A) is already bound?

In that case (see Methods section for details), the function for E_LAB_ changes to

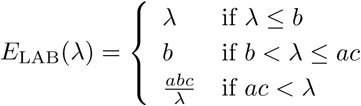

How is this cooperative case different from the non-cooperative case? It is easy to see that the maximum number of bound complexes is still the same, because this is determined by b (in other words, the availability of the scarcer of the two ligands). Two things, however change: First, the range of concentrations at which this maximum number of complexes is formed, becomes larger, i.e. we can increase *λ* further without seeing a detrimental effect on LAB formation. Second, after the maximum is reached, the decline in the expected number of LAB complexes as a function of *λ* is less steep. There is still a hook effect, but the effect is less drastic, and it sets in at higher concentrations of L. This is how cooperative binding works to counteract the hook effect. Figure 4 shows the cooperative case for the same values of *a*, *b*, and *λ* as the non-cooperative example shown above.

**Figure 4.**
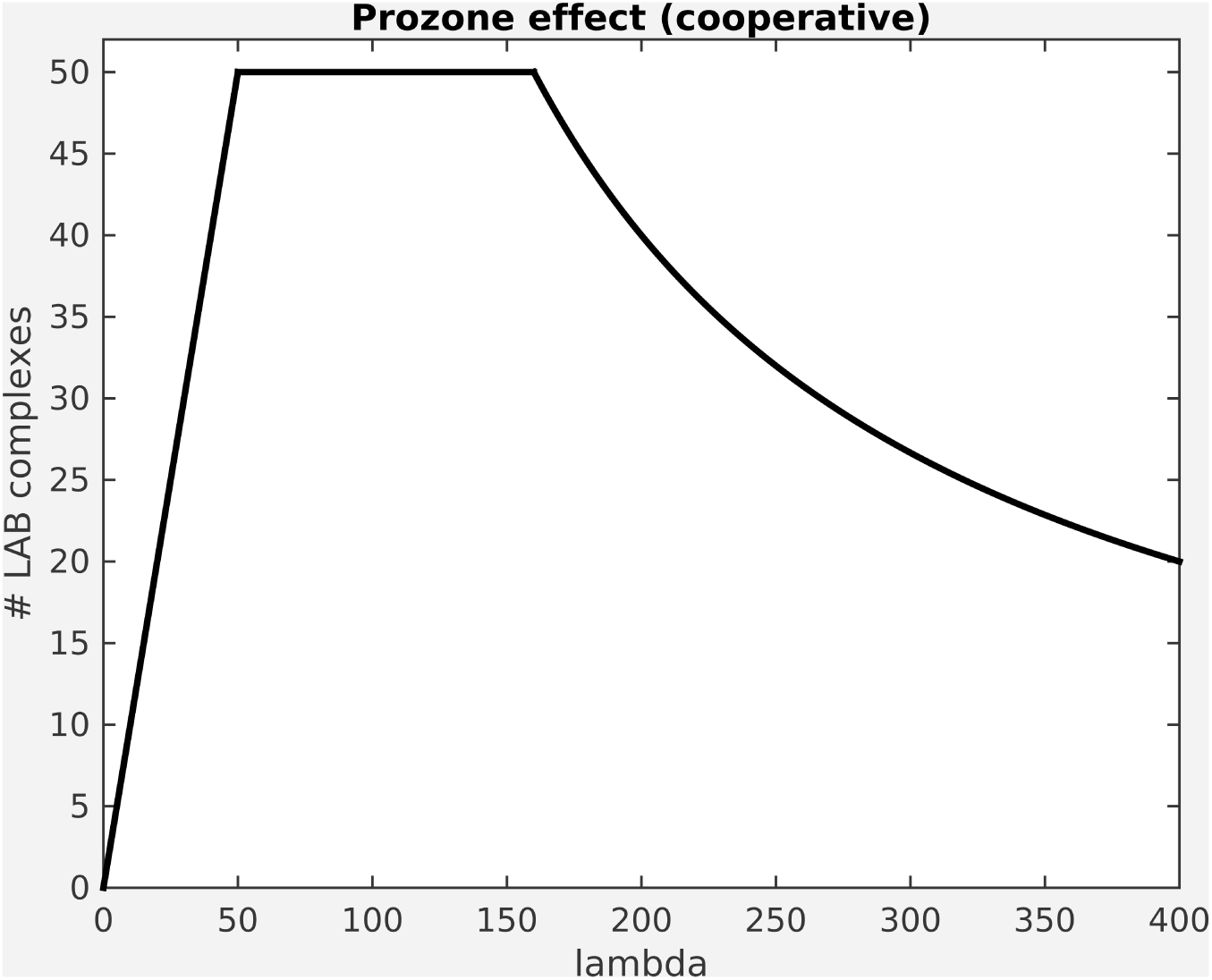
The Prozone effect for a Linker protein with cooperativity, assuming perfect binding. Expected values are shown. Amount of linker protein (lambda) varied from 1 to 400, amounts of proteins A and B were 80 and 50, respectively. The cooperativity constant c was set to 2. Plot was drawn in MATLAB [36].

The above analysis assumes that binding of A and B to L is perfect, in the sense that if there is a free molecule of ligand and there is an unoccupied binding site, then binding will happen with a probability of 1. In real biological systems, of course, such certainty does not exist. The probability of a binding event depends not only on the availability of ligand and binding sites, but also on their affinities, usually measured in terms of association or dissociation constants.

This will affect the expected number of fully bound complexes, the range of concentrations at which certain behaviours can be observed, and the way we think about cooperativity. An analytical analysis is complicated by the fact that, unlike in most other binding scenarios that are well described in theoretical biochemistry, we are operating under conditions of “ligand depletion”, where the limited availability of ligand will affect the dynamic behaviour of the system [12].

Therefore, the scenario of real-life biological systems with non-zero dissociation constants lends itself well to simulation approaches. In simulations of biochemical systems, one possible way of representing cooperative binding is as a decrease in dissociation constants (i.e. an increase in affinity) if one or more of the binding sites on the receptor are already occupied [10].

### Calmodulin binding to calcium displays a high-dose hook effect

In order to investigate whether we can detect a hook effect in a simple linker protein under conditions found in biochemical systems (with finite association constants), we examined the high-dose hook effect using an earlier model of calmodulin activation by calcium [13].

Calmodulin is a calcium-sensing protein that has an important role in bidirectional neuronal plasticity. In the post-synaptic neuron, it acts as a “switch” between induction of long-term potentiation (LTP) and long-term depression (LTD), by activating either Ca^2+^-/calmodulin-dependent kinase II (CaMKII) or calcineurin, respectively (reviewed in [14]). The decision to activate either one or the other depends on the input frequency, duration and amplitude of the postsynaptic calcium signal [11]. Each calmodulin molecule binds to four calcium ions in a cooperative manner [15]. Structural evidence [16, 17] suggests that this cooperativity arises from allosteric regulation. According to this model [18, 13], calmodulin can exist either in the T state with lower calcium binding affinities or in the R state with higher calcium binding affinities. The more calcium ions are bound to a calmodulin molecule, the higher the likelihood that it will transition from the T state to the R state.

Other models of calmodulin regulation exist [19, 20], but for our purposes of examining the relationship between cooperativity and the hook effect, the allosteric model proposed by Stefan et al. [13] is sufficiently detailed. The model accounts for two states of calmodulin (R and T) and four calcium binding sites, with different calcium affinities. In addition, R state calmodulin can bind to two allosteric activators, CaMKII or calcineurin (PP2B).

As expected, wildtype calmodulin displays a high-dose hook effect, as shown in the black line in figure 5: If we plot the formation of fully-bound calmodulin (calmCa4) as a function of the initial calmodulin concentration, then the curve initially rises, but then drops again at high doses of calmodulin, indicating that calmodulin molecules compete with each other for calcium binding.

**Figure 5.**
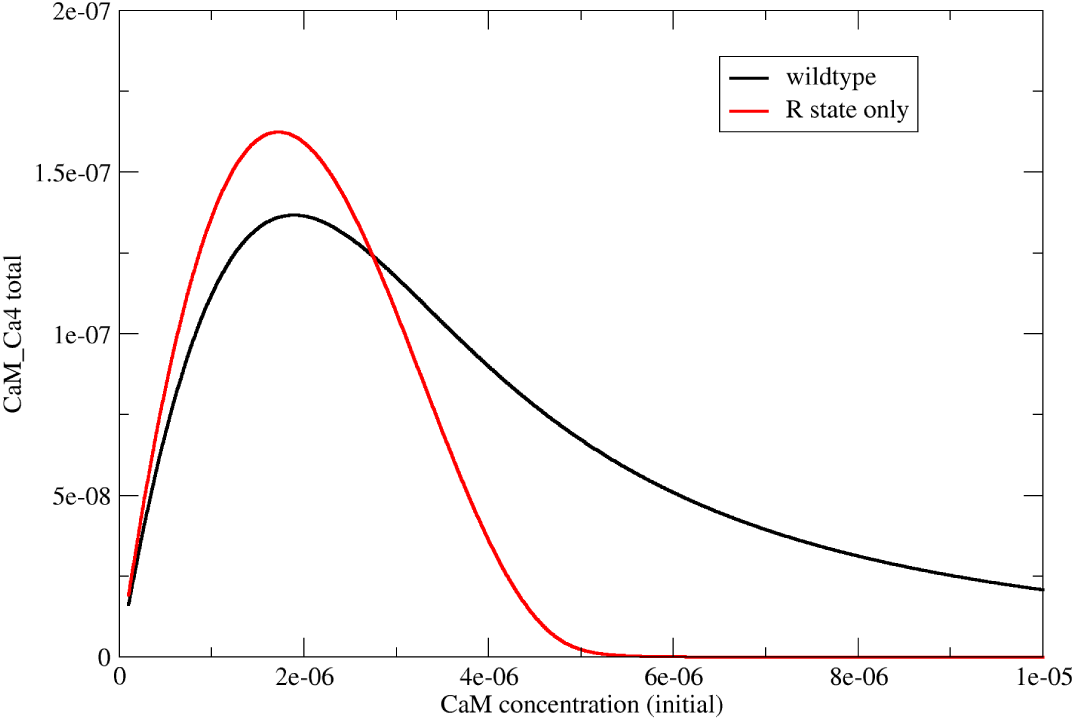
Reduced hook effect in cooperative (wt) calmodulin. This figure shows the results of simulations on wildtype calmodulin (which is allosterically regulated, in black) compared to a non-cooperative *in silico* mutant (R state only, in red). The plot of fully bound calmodulin as a function of initial calmodulin concentration shows a prozone effect in both cases, but it is more pronounced in the non-cooperative version.

Is the high-dose hook effect dependent on our particular parameter choices? In this model, we used parameters for dissociation constants and R-to-T transition that had previously been shown to produce simulation results consistent with the available literature on calmodulin binding to Calcium under a variety of conditions [13]. Nonetheless, we repeated the simulations at varying dissociation constants and varying values of *L* (which governs the transition between R and T states). As shown in Additional file 3, a high-dose hook effect exists in a variety of parameter regimes, although it can be more or less pronounced.

### Allostery mitigates the high-dose hook effect in calmdoulin

If it is true that cooperativity helps mitigate the prozone effect, then a non-cooperative protein with similar properties to calmodulin would show a higher hook effect than calmodulin itself. To test this hypothesis, we created an artificial *in silico* variant for calmodulin that binds to calcium in a non-cooperative way. This was done by abolishing R to T state transitions in the model, so that calmodulin could exist in the R state only. It is important at this point to differentiate between affinity and cooperativity: The R state only version of calmodulin has higher calcium affinity than the “wildtype” version (which can exist in the R state or the T state). But the R state only version has itself no cooperativity, because cooperativity arises from the possibility of transitioning between the T and R states [21, 22].

Figure 5 shows the results of two simulations run on wildtype calmodulin and an R-state-only *in silico* mutant, respectively. Plotting fully bound calmodulin as a function of calcium concentration reveals a high-dose hook effect in both cases. However, despite the R-state only variant reaching a higher peak (due to its higher overall affinity), it also shows a more pronounced hook effect, with lower absolute levels of fully bound complex at higher calmodulin concentrations.

### Molecular environment modulates calmodulin cooperativity and hence, susceptibility to the high-dose hook effect

We have shown that calmodulin binding to calcium can be affected by the hook effect, and that this hook effect is stronger in non-cooperative versions of calmodulin. In order to assess the relevance of these findings for the cellular function fo calmodulin, we need to answer two questions: First, are the concentration regimes under which this system displays a hook effect ever found under physiological conditions? And second, are there existing forms of calmodulin that resemble our “R state only” or “T state only” *in silico* mutations and are therefore non-cooperative? Calmodulin is found in various concentrations in various tissues of the body, from micromolar concentrations in erythrocytes to tens of micromolar concentrations in some areas of the brain [23]. The calmodulin concentrations used in our simulations are therefore physiologically relevant, especially in the higher range, where the prozone effect is most pronounced.

Our mathematical treatment and simulations have shown that allosteric regulation mitigates the hook effect. But what is the relevance of this for calmodulin? After all, there is no known variant of calmodulin that exists only in the R state or only in the T state. However, there are allosteric modulators that will stabilise one of the two states, and they can exist in high concentrations. To investigate the effect of the presence of an allosteric modulator, we repeated the above simulations in the presence of 140 *μ*M CaMKII. This number is consistent with the number of CaMKII holoenzymes found in post-synaptic densities in labelling studies [24].

The results of our simulations in the presence of 140 *μ*M CaMKII are shown in figure 6 a. Since CaMKII is an allosteric activator, it stabilises the R state of calmodulin over the T state. At such high concentrations of CaMKII, the R state dominates, and calmodulin behaves almost like the theoretical R-state-only form. In particular, the hook effect is exacerbated at high calmodulin concentrations.

**Figure 6.**
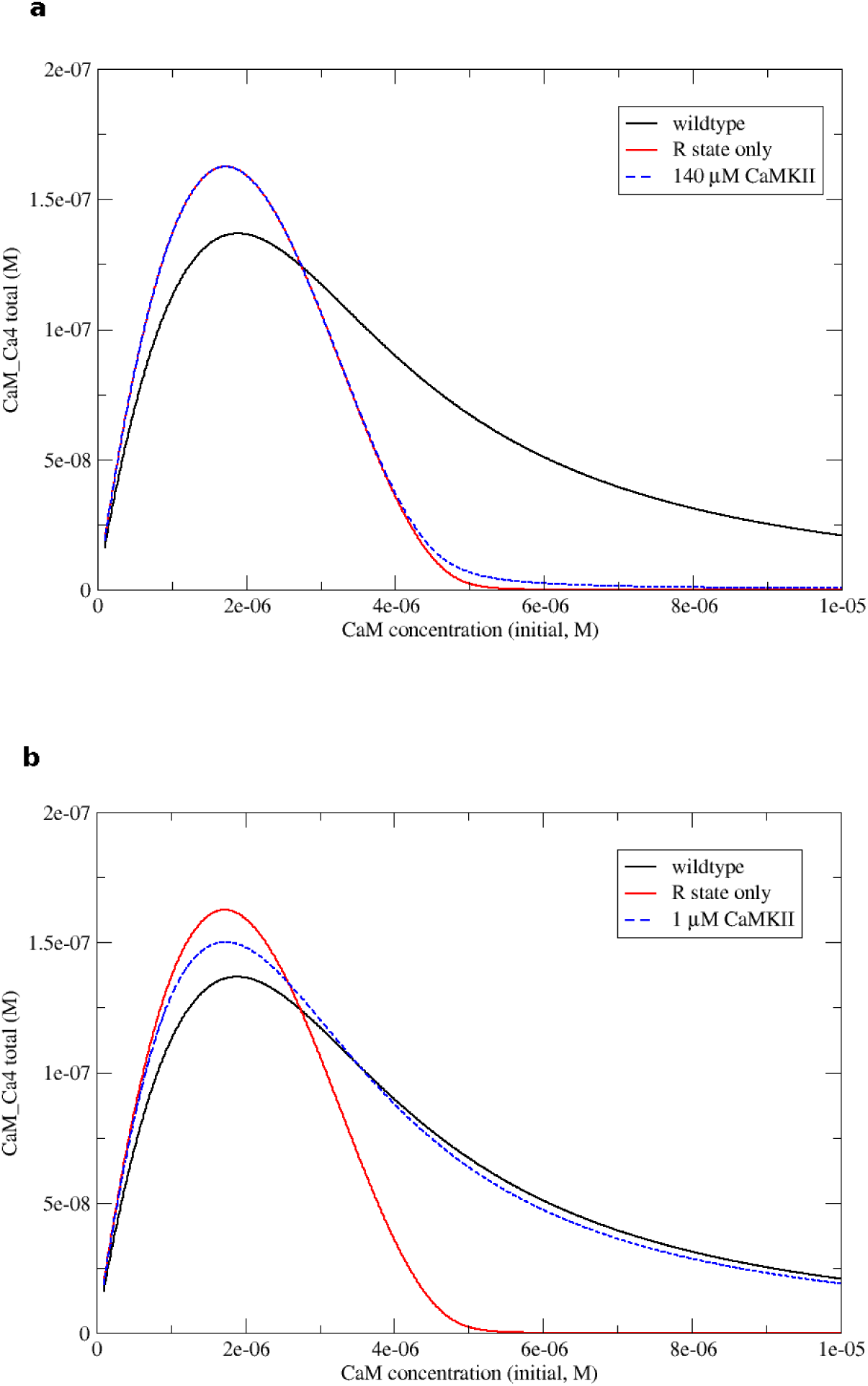
Allosteric modulators can exacerbate the hook effect by reducing cooperativity. As in the previous figure, we show fully bound calmodulin as a function of initial calmodulin concentration, both for wildtype calmodulin (black) and an *in silico* mutant that exists only in the R state (red). We also show the results of adding two concentrations of the allosteric activator CaMKII (blue). **a** At 140 *μ* M CaMKII, calmodulin exists almost exclusively in the R state and thus behaves like the non-cooperative *in silico* mutant. **b** at 1 *μ* M CaMKII, both states exist and the prozone effect is comparable to wildtype calmodulin.

To assess the effect of concentration of the allosteric activator, we compared this scenario with one where the CaMKII concentration was reduced to 1, *μ*M. In this case (shown in figure 6 b) the R state is stabilised to some extent, but R and T states still co-exist, and cooperativity is therefore preserved. While the initial peak of fully bound complex is higher than for wildtype calmodulin in the absence of any allosteric effectors, the prozone effect is reduced.

Taken together, this indicates that under conditions that render a protein susceptible to the high-dose hook effect, higher concentrations of an allosteric activator result in less activity than lower concentrations.

## Discussion

### Cooperativity gives partially bound linkers a competitive edge

In this study we asked whether there is a general principle by which allosterically regulated proteins such as calmodulin are - to some extent - protected from the high-dose hook effect. To mathematically examine this question, we have used combinatorics to show how the high-dose hook effect arises in a simple trimolecular complex with perfect binding affinities. This is, in essence, due to the linker protein competing with other instances of itself for full complex formation. This result reproduces the one found by Ha and colleagues, who derived algebraic expressions for all concentrations in a similar system and systematically varied dissociation constants and concentrations of components to explore the prozone effect [8].

In addition, we also show that cooperative binding mitigates the high-dose hook effect. It does so by essentially giving partially bound versions of the linker protein a competitive advantage, so that the population is skewed towards either fully bound forms or fully unbound forms, at the expense of partially bound forms.

### Cooperativity can protect calmodulin from the high-dose hook effect under physiological conditions

Calcium binding to calmodulin is cooperative, and this suggests that calmodulin would be protected, to some extent, from the high-dose hook effect. Indeed, we could show that is the case for physiological ranges of calmodulin concentration. However, the cooperative nature of calmodulin binding to calcium itself is not a fixed property, but can vary according to the cellular environment. Cooperativity can be reduced under conditions of ligand depletion [12], which are also the concentration regimes where the hook effect becomes noticeable. In addition, high concentrations of an allosteric modulator can reduce cooperativity. Thus, the susceptibility of a protein to the high-dose hook effect depends not only on intrinsic properties of the protein and its ligand, but also on the cellular context.

### More is not always better

As we have seen, the presence of an allosteric activator can reduce cooperativity. This is because cooperativity in allosteric molecules arises, fundamentally from the ability of the molecule to transition between T and R states, which have different ligand affinities. By pulling all of the allosteric protein towards either the T or the R state, cooperativity is reduced, and the high-dose hook effect becomes more pronounced. Interestingly, this is true no matter whether it is the T state or the R state that is stabilised or, in other words, whether the allosteric modulator is an inhibitor or an activator. Thus, under conditions where the hook effect is noticeable, allosteric activation behaves counterintuitively: There is less activity in the absence of the allosteric activator than in its presence, and less activity when the levels of allosteric activator are high than when they are low.

### Possible experimental validation

Our model predicts that the presence of cooperativity protects, to some extent, against the high-dose hook effect.

Ha and Ferrell [25] have investigated the link between cooperativity and the highdose hook effect using a binding system composed fo three DNA strands: One strand can bind to two others, and cooperativity can be engineered by tweaking with the amount of overlap. Indeed, the construct identified as positively cooperative showed a less pronounced hook effect than the construct identified as non-cooperative [25]. This corroborates our ideas in a synthetic binding system.

In order to assess whether this is also the case for systems with more than two binding sites, and in particular for multivalent proteins, one would need to be able to do things: First, to be able to measure full occupancy of some multivalent protein (without measuring partially occupied states). Second, this would need to be done on a linker protein of which there are two related forms, one of which shows cooperative binding and the other does not.

For calmodulin, as we have seen, a non-cooperative (or less cooperative) state can be obtained by adding a large concentration of CaMKII. This will stabilise the R state, which is not itself cooperative. In contrast, in the absence of allosteric modulators, both the R and T states are populated, and the transition between them is what confers cooperativity to calmodulin. Thus, creating a less cooperative form of calmodulin in vitro is easy. However, measuring full saturation of calmodulin is not. Fractional saturation (i.e. the ratio of occupied binding sites) cannot serve as a proxy, because it does not show a Hook effect, instead monotonically going down as calmodulin concentration increases, as expected. In addition, fractional occupancy profiles in the absence and presence of CaMKII do not show a big difference (see Additional File 3). Thus, fractional occupancy is not a good proxy for full saturation. Instead, is it possible to measure conformational state? Moree et al. have shown that it is possible to measure conformational change in calmodulin that occurs with Calcium binding [26]. However, conformational state and full saturation do not directly translate into each other, as can be seen in Additional File 3, where we plotted *R̄* for calmodulin in the absence and presence of CaMKII. Thus, measuring fully saturated calmodulin (and therefore assessing the magnitude of the high-dose hook effect *in vitro*) is non-trivial. As molecular measurement techniques develop in the coming years, though, this work provides a hypothesis that will be amenable to testing.

Another possibility would be to compare hemoglobin and myoglobin. Both have similar properties, but hemoglobin exists as tetramer exhibiting cooperative binding to oxygen, while myoglobin is monomeric and therefore non-cooperative (reviewed in [22]). Obviously, since myoglobin has only one oxygen-binding site, it does not itself display a high-dose hook effect. Instead, it could be used as a proxy for what a non-cooperative version of hemoglobin would look like. The fraction of fully occupied “non-cooperative hemoglobin” can simply be computed by taking the fractional saturation of myoglobin to the fourth power (essentially grouping free myoglobin molecules into groups of four and declaring full occupation if all four are bound). The prediction is that this would show a stronger high-dose hook effect than hemoglobin itself.

### Relevance to other systems

These results are likely to be relevant in a wide range of biological systems. For instance, neuronal signalling depends on a number of proteins with multiple ligand binding sites, including membrane receptors such as the AMPA receptors, NMDA receptors or other postsynaptic calcium sensors such as calbindin. The existence of multiple ligand binding sites and, under some conditions, the relative scarcity of ligands (e.g. of glutamate in the synaptic cleft, and of calcium in the postsynaptic neuron) makes those proteins, in principle, prone to the hook effect. Interestingly, several of these proteins are allosterically regulated (this is the case, for instance, for AMPA receptors[27] and for NMDA receptors[28]), which could confer a sensitivity advantage at high receptor-to-ligand ratios [12].

The hook effect is also a frequently discussed problem in medical diagnostics, because it can lead to false-negative effects if the levels of analyte to be detected are too high. Recent examples of this effect have been reported in the diagnosis of meningitis[29], malaria[30, 31], and even in pregnancy tests[32]. To avoid such cases, systematic dilution of the sample (and thus a reduction of analyte concentration) can help[33], but is not always practicable[34]. Given our results, another way to reduce the risk of false-negative results due to the hook effect would be to somehow make analyte binding to the reporter in the assay cooperative. One way of achieving this in a sandwich immunoassay by making one of the receptors multimeric has been patented in 2001[34].

## Conclusions

If a protein acts as a linker between different parts of a multimolecular complex, then there are concentration regimes where adding more of the linker protein to the mixture will result in less overall complex formation. This phenomenon is called the high-dose hook effect or prozone effect.

We have provided an idealised mathematical description of the hook effect and shown that allosteric regulation does indeed mitigate the hook effect, as has been predicted before [9].

Whilst this means that allosteric proteins such as calmodulin are, to some extent, protected from the high-dose hook effect, the presence of allosteric modulators can increase susceptibility to the high-dose hook effect. The extent of the hook effect is therefore strongly dependent on the cellular microenvironment.

## Methods

### Complex formation curve for LAB

Assume a perfect binding system with *λ* molecules of a linker molecule L, where every molecule of L can bind to one molecule of A and one molecule of B. Numbers of A and B are denoted by *a* and *b*, respectively, with *b* ≤ *a* (wlog).

Assuming perfect binding and no cooperativity, the molecules of A and B will be distributed randomly across molecules of L. At the end of the binding phase, any given molecule of L will be either free, bound to A only, bound to B only, or part of a complete LAB complex. Clearly, this is a combinatorial problem that can have a variety of possible outcomes in terms of the numbers of complete LAB complex, partial complexes (LA or LB) and free (unbound) L.

We are interested in expressing the expected number of full complexes (LAB) formed as a function of *λ*. We will denote this quantity as *E*_LAB_(*λ*)

As long as the number of linker proteins L is limiting, then the total number of ternary complexes formed will be *λ*.

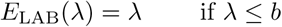

If the amount of linker protein is larger than the amount of protein B, but smaller than the amount of protein A, then all of L will be bound to A at least, and the amount of completely formed LAB complex will depend on b alone.

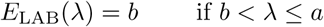

Finally, if the amount of linker protein is larger than both *a* and *b*, then we have to consider all possible binding scenarios. Figure 3 shows a probability tree for each molecule of L (with *c* = 1 in the absence of cooperativity). For reasons of convenience, we show binding as a two-stage process (A binds first, then B), but this is not meant to represent a temporal order. The resulting probabilities for each end state would be the same if the order of binding was switched.

The expected number of LAB complex can be computed by taking the probability of each L to become an LAB complex, and multiplying with the amount of L:

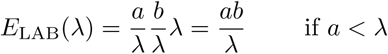

Thus, for fixed amounts of A and B (with *b* ≤ *a*), we can write the expected amount of LAB as a function of *λ* as a three-part function:

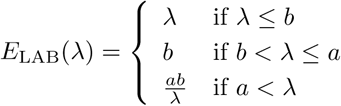

### Formation of LAB complex if ligand binding is cooperative

The case where ligand binding is cooperative (i.e. where binding of a molecule of A facilitates the binding of a molecule of B to L, and vice versa) is analogous.

Again, as long as *λ* is smaller than both *a* and *b*, the amount of linker L will be limiting, and we thus have:

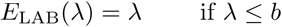

If the amount of linker protein is larger than the amount of protein B, then there can be at most *b* fully bound complexes, just like in the non-cooperative case. Thus, *b* is the maximum possible value for E_LAB_.

If *λ* exceeds both *a* and *b* by a sufficient amount, we can again follow a probability tree (displayed in figure 3) to determine the probability of a single linker protein being fully bound. Again, this is computed as the probability of A binding (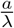, as before) times the probability of B binding, given A is already bound, which will depend both on 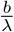 (as before)and on a cooperativity coefficient *c*. This is a coefficient that modulates the probability of subsequent binding events, with *c* > 1 indicating positive cooperativity and *c* = 1 no cooperativity. For instance, for calmodulin binding to calcium, *c* would be around 3 and for hemoglobin binding to oxygen around 1.5 (computed from dissociation constants reported in [35]). This gives us an expected value for the number of fully formed LAB complexes:

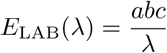

What do we mean by “a sufficient amount”? Clearly, *λ* must be bigger than both *a* and *b*. But remember also that E_LAB_ is limited by *b*. So, the question is, when is 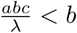? This is the case when *ac < λ*.

Thus, the complete function for E_LAB_ is as follows:

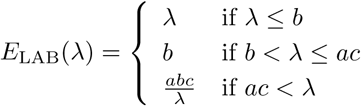

### Theoretical complex formation curves

The complex formation curves under the assumption of perfect binding shown in figure 2 were generated using MATLAB [36]. We also used MATLAB to simulate 100 cases of A and B binding to L as follows: For each simulation step, *a* (or *λ*, if *λ < a* binding sites were randomly chosen and defined as bound to A, *b* (or *λ* if *λ < b*) binding sites were chosen and defined as bound to B. The number of binding sites occupied by both A and B was then determined and plotted. The MATLAB script used to generate the plots is provided as Additional File 1.

### Calmodulin simulation

For simulations of the prozone effect in calcium binding to calmodulin, we used a model of calmodulin published earlier [13]. The model accounts for two conformational states of calmodulin (R and T) and four different calcium binding sites (A, B, C, D). In addition, R state calmodulin can bind to CaMKII or PP2B. The full model is available in BioModels Database [37] as BIOMD0000000183.

For simulations of calcium binding to wildtype calmodulin, the concentrations of both CaMKII and PP2B were set to zero.

For simulations of the R state (or T state) only, the transition rates between R and T state were set to zero, and the initial concentration of calmodulin was set to be all in the R (or T) state. For simulations in the presence of an allosteric activator, we used a CaMKII concentration of 140 *μ*M, which corresponds to reports of typical levels of around 30 holoenzymes of CaMKII found in post-synaptic densities with a volume of around 5 × 10^−18^ *l*[24]. To test the effect of reducing CaMKII concentration, simulations were run again setting CaMKII concentration to 1 *μ*M.

Simulations were run using Copasi[38]. The simulations took the form of a parameter scan over initial calmodulin concentrations ranging from 10^−7^ to 10^−5^ M in 1000 steps. The scan was over free calmodulin T for all simulations, except for the “R state only model”, where the scan was over free calmodulin R. All other calmodulin species were initially set to 0. Each parameter scan simulation was a time course lasting 1000 seconds, which was in all cases largely sufficient to equilibrate the model.

All simulation results were plotted in Grace (http://plasma-gate.weizmann.ac.il/Grace/).

## Declarations

## Ethics approval and consent to participate

Not applicable.

## Consent for publication

Not applicable.

## Availability of data and materials

The calmodulin model is based on an earlier calmodulin model by Stefan et al[13], which is available on BioModels Database under the following link: https://www.ebi.ac.uk/biomodels-main/BIOMD0000000183. The version used here differs in the concentrations of molecular species involved; the model with calmodulin only is included as a supplementary material to this article; the “R only” and “T only” mutants can be generated by setting the parameter kRT to zero and adjusting the initial calmodulin concentration. Simulations in the presence of CaMKII can be generated by setting the initial CaMKII concentration as needed. Matlab code used to generate the figures in the first part of the paper is also available in the supporting information.

## Author’s contributions

Developed theoretical framework: MIS; designed and performed simulations: RDR,MIS; analysed and discussed results: RDR, CR, MIS; wrote the paper: RDR, MIS.

## Acknowledgements

The authors thank members of the Le Novère Lab at the Babraham Institute, Cambridge (UK) for helpful discussions.

## Additional Files

Additional file 1 — Matlab code used to produce figures 4 and 6

Additional file 2 — Copasi file for the calmdoulin model

Additional file 3 — Supplemental text

